# Perceiving latent dynamics: Innate and coachable visual estimation of limb damping

**DOI:** 10.64898/2026.03.02.708982

**Authors:** Taliyah Huang, Meghan E. Huber, Jeremy D. Brown, A. Michael West

## Abstract

Humans are remarkably adept at extracting latent dynamic information from purely visual cues. Prior work shows that people can innately estimate differences in limb stiffness using solely their visual observation of movement, which suggests that components of mechanical impedance may be embedded within humans’ internal predictive models of movement. We tested whether humans can similarly perceive damping, a force-velocity relationship, and whether targeted coaching can enhance this visual ability. Specifically, 30 participants observed abstract two-link arm simulations with systematically varied elbow damping and rated their perceived level of damping for several trials. Participants completed two sessions separated by one of three brief coaching interventions: (1) no coaching, (2) coaching to attend to hand velocity, or (3) coaching to attend to elbow-angle velocity. Results reveal that (1) humans can innately perceive changes in arm damping using solely their visual observation of motion and (2) coaching further improved performance, with the elbow-angle coaching group showing a significantly greater increase in rating accuracy compared to the other two groups. This work extends our understanding of how action-perception coupling supports inference of mechanical impedance. Moreover, we demonstrated that perceptual strategies for estimating damping are malleable and can be systematically improved through coaching. We not only identified the visual cues observers relied on but also guided them toward more classifiable features, effectively strengthening their perceptual models of limb dynamics.

**Author summary:** Humans are remarkably adept at understanding an object’s latent dynamic properties simply by watching it move, even when the underlying forces are unseen. In this paper, we demonstrated that people can notice differences in how “damped” a moving limb is using vision alone. Moreover, we found that brief coaching helped participants focus on the most informative features, significantly improving their ability to differentiate the damping levels. These results demonstrate how people can visually infer aspects of movement that are normally thought to require physical interaction, offering insight into how the motor system links action and perception. They also show that strategies can be shaped and improved, supporting real-world healthcare applications. In stroke rehabilitation, physical therapists physically assess the resistance of a patient’s limb, so better guidance on the most relevant visual cues can help clinicians learn faster and even provide care remotely. In robot-assisted surgery, surgeons operate a console to perform procedures with limited or no force feedback, so they must estimate tissue dynamics properties largely from visual observation. Understanding how people visually estimate these dynamics can inform training for more precise surgical decisions. Overall, our findings clarify how humans interpret movement dynamics and how coaching can support more consistent and accurate perceptual decisions.

## Introduction

Humans can remarkably interpret the movement dynamics of other people. Classic demonstrations of biological motion perception show that people can identify familiar figures, recognize gait patterns, and even infer intent or emotion from sparse point-light displays [1–4]. These findings highlight that visual observation alone can provide access to *latent* properties of movement that are not directly observable in the kinematics. In many cases, observers appear capable of recovering aspects of the underlying physical interactions that generate a motion pattern, even when the forces and neuromotor commands are hidden from view. This ability suggests that humans treat movement as a structured signal that encodes deeper dynamical information, motivating the question of which mechanical properties are visually accessible and how the motor system supports their estimation.

Prior work has shown that humans can extract not only kinematic information from observed motion but also latent dynamic properties. In particular, recent evidence demonstrates that people can visually estimate differences in limb stiffness using only the motion of a simulated arm [5–7]. This finding is surprising because stiffness is formally defined as a force to displacement relationship, as described by Hooke’s law *F* = *kx*, and therefore cannot be recovered from kinematics alone [8]. Observers in these tasks received no information about the forces that generated the observed motion, yet they produced reliable stiffness judgments. This suggests that people may rely on internal predictive models that link observed movement patterns to the impedance properties that could have produced them.

A growing body of research suggests that the visual interpretation of movement is closely tied to the motor system that produces it. Neuroimaging and neurophysiological studies indicate that many of the brain regions engaged during action execution are also recruited during action observation [9–11]. This shared circuitry is often associated with mechanisms that allow observers to internally simulate the movements they observe, effectively linking perceived kinematics to the types of motor commands that could have produced them. Complementary behavioral work shows that humans can recognize, imitate, and learn motor behaviors through observation alone [1, 12, 13]. These findings support the broader idea that the visual system does not treat motion as a purely geometric signal but instead interprets it through internal models shaped by one’s own sensorimotor experience. Because the regulation of mechanical impedance is a central principle of how humans control and stabilize movement [14–18], this framework raises the possibility that observers may also access other impedance-related properties, such as damping, through visual motion.

Damping describes the resistive forces that oppose motion in proportion to velocity, captured by the relation *F* = *bv*, and therefore cannot be directly recovered from kinematics without access to the underlying forces that shape the trajectory. Like stiffness, damping is a latent physical property that influences how a limb moves through space, yet its effects are expressed only indirectly through the resulting pattern of motion. Establishing whether damping is visually accessible would clarify whether humans can infer multiple components of impedance through observation and deepen our understanding of how the perceptual system interprets dynamics from movement. As such, in this paper, we assess whether humans can estimate changes in damping using solely the visual observation of motion.

If damping is visually accessible, observers must rely on specific motion features to infer it. This raises the question of whether these visual strategies are innate or can be learned and tuned. A large body of work on perceptual learning shows that repeated experience, targeted instruction, and attentional guidance can substantially improve visual discrimination, including sensitivity to complex or subtle motion cues [19–24]. Similar principles appear in motor learning, where observation and feedback contribute to the refinement of internal models and the acquisition of new skills [25]. Perceptual training can also enhance performance in demanding real-world tasks, such as ultrasound image interpretation, which further suggests that visual strategies can be shaped by structured experience [26]. These findings raise the possibility that coaching can help observers (1) shift their attentional strategies toward the motion features that are most informative for estimating damping and (2) strengthen the internal representations that support this perceptual ability, ultimately improving estimation performance. Thus, in this study, we assess whether the perception of damping can be improved through targeted coaching.

These possibilities are relevant not only for understanding basic perception–motor processes but also for contexts in which people routinely judge limb dynamics, such as in the clinical evaluation of rigidity and spasticity. Rigidity and spasticity are displacement- and velocity-dependent abnormalities in muscle tone that significantly affect the quality of life of stroke survivors and are central markers of motor impairment [27]. In current clinical practice, physical therapists assess these impairments through hands-on examinations where they judge resistance during passive limb movement. Since this assessment relies on subjective impressions, it can vary widely across clinicians [28]. Importantly, rigidity and spasticity map directly onto the mechanical properties of stiffness and damping, respectively, making them the clinical analogs of the latent dynamical properties investigated in this line of research. Understanding whether humans can visually estimate these properties from movement alone may reveal the motion features that clinicians implicitly rely on when evaluating limb impedance. It may also motivate more quantitative and reliable approaches to assessing these impairments.

Together, these considerations motivate a controlled investigation of how well humans can visually estimate damping and whether this ability can be improved through targeted coaching. Accordingly, we designed the present study to address the following research questions:

- **RQ1:** Can humans innately estimate differences in limb damping from visual observation alone when observing movements generated by a simulated two-link arm?
- **RQ2:** Can coaching change the visual attention strategies observers use when estimating damping?
- **RQ3:** Does targeted coaching improve damping-perception accuracy by guiding observers toward informative motion features?

Addressing these questions will deepen our understanding of how humans interpret the dynamical structure of movement from vision alone and clarify *if* and *how* the perceptual models that support this ability can be enhanced through coaching.

## Materials and Methods

### Participants

A total of 30 participants (15 males and 15 females with a mean age of 24.03±4.79 years) took part in the experiment. Participants had a variety of educational backgrounds, and none had any prior experience with the experimental task. All participants gave informed written consent before the experiment. The experimental protocol was reviewed and approved by the Institutional Review Board of the Johns Hopkins University School of Medicine (IRB Protocol #00263386).

### Experimental protocol

While sitting in front of a computer screen, participants observed a simulation of a two-link arm with an undisclosed level of joint damping and were asked to rate the perceived damping of the arm (Fig. 1). Specifically, the participant would view the 2D simulation for 25 seconds or click on the “skip” button that appeared on the screen after 6 seconds, which would take them to the rating screen allowing them to submit their perceived damping on a scale of 1 (least damped) to 7 (most damped). Then, the participant would be able to self-initiate the next trial. For an example, see S3.

**Fig 1.**
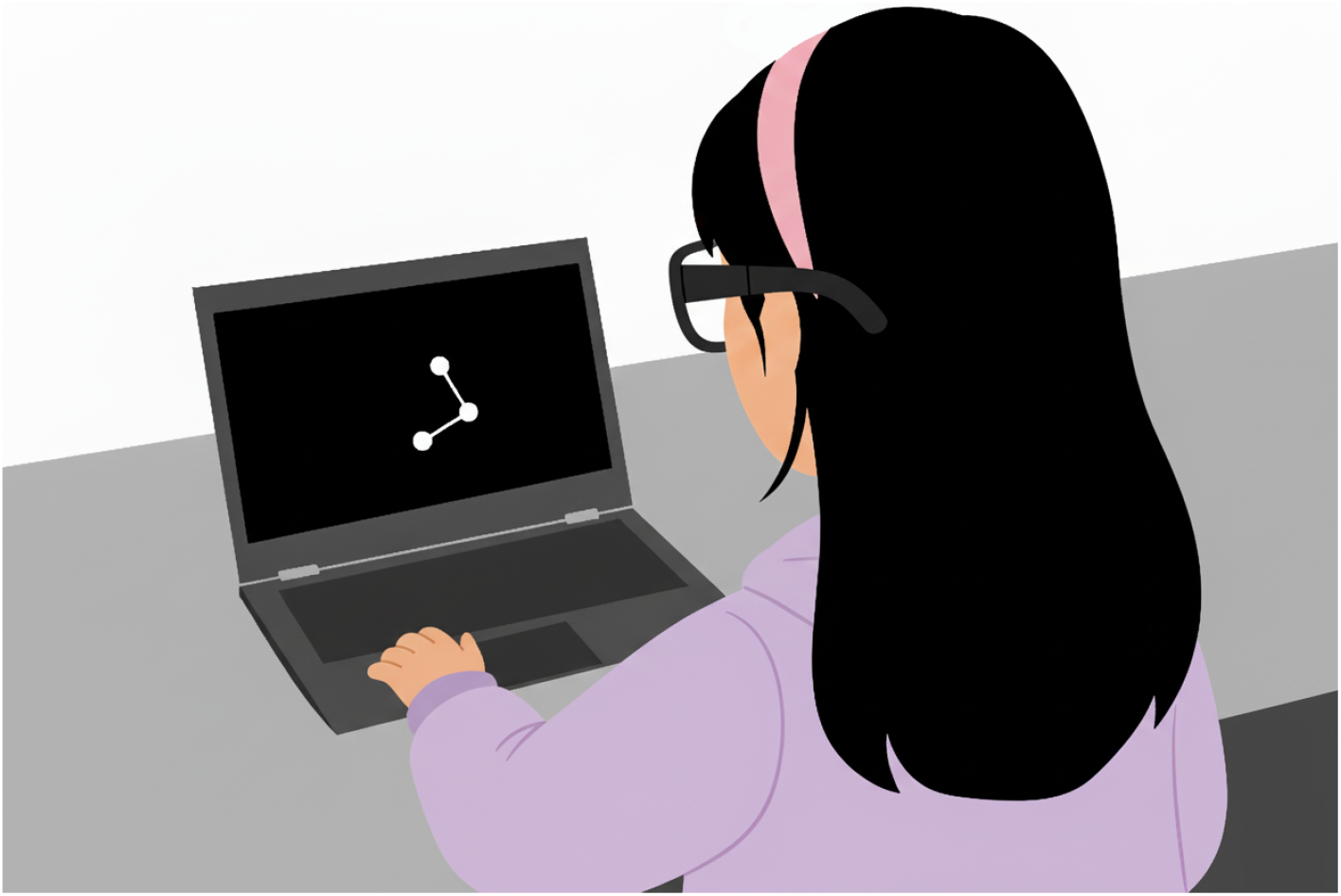
Experimental setup. In each trial, participants were instructed to observe the motion of an abstract 2D display of a two-link planar arm move and then estimate its damping on a numeric scale from 1 (“least damped”) to 7 (“most damped”). This figure was generated with ChatGPT and Gemini.

The experiment was divided into two sessions with a coaching intervention in between (described in detail below). In each session, each participant viewed 30 arm simulations with six predefined damping levels at the elbow joint (0, 2, 4, 6, 8, 10 Nms/rad) in a block-randomized order (five blocks, each consisting of six randomized damping levels). Participants were not provided with any details about the observed simulations.

After each session, participants were asked to fill out a digital survey. First, they provided a subjective rating from 1 (lowest) to 7 (highest) of the following standard NASA Task Load Index (TLX) metrics [29]: Mental demand, physical demand, temporal demand, effort, frustration, and performance.

Next, the participants were asked about their strategy with a simple free response question (“Describe your strategy when making your ratings for this past block of 30 simulations.”). To acquire more quantifiable information about strategy, participants were also asked to (1) estimate the percentage of time they believed they were looking at specific *locations* on the two-link arm simulation (i.e., hand, elbow, angle, other; Fig. 2) and (2) estimate the percentage of time they believed they were analyzing various *motion* characteristics (i.e., displacement, velocity, acceleration, jerk, other).

**Fig 2.**
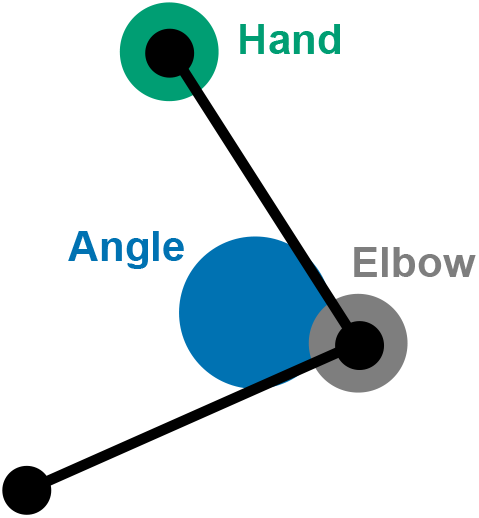
Locations diagram on the survey. Participants were shown this diagram in the self-report survey as they estimated the percentage of time they spent looking at each of the labeled positions.

### Eye-tracking

Furthermore, to provide quantitative validation of the location strategies described by participants in the self-reported survey, participants wore eye-tracking glasses (Pupil Core, Berlin) to monitor their gaze as they watched the simulations. However, due to calibration and camera-registration issues, the majority of the eye-tracking data was too noisy to analyze. Thus, we moved the discussion of the eye-tracking to Supporting Information.

### Coaching

In between the first and second sessions of 30 trials, participants were given a coaching intervention. Each participant was randomly assigned to one of three coaching groups: “No Coaching,” “Hand Coaching,” and “Angle Coaching.” Each coaching group consisted of 10 different participants. The No Coaching group received no coaching intervention and moved on to Session 2 after a *∼*2 minute break. Participants in the Hand Coaching group watched a 2-minute “Hand Coaching video” between Session 1 and Session 2 that explained the strategy of observing the varying *hand speed* as it traveled in a loop to estimate damping (See S1). Participants in the Angle Coaching group watched a 2-minute “angle coaching video” that explained the strategy of observing the elbow *angle speed* (rate of the arm’s opening and closing) to estimate damping (See S2). Both videos highlighted that “the slower the speed, the higher the damping.”

Following the coaching intervention, the second session was conducted identically to the first, with participants viewing and rating an additional 30 randomized damping simulations while wearing the eye-tracking glasses, and then completing the same surveys as described above.

### Simulations

To generate the arm motions shown to participants, we developed a custom MATLAB program that produced two-link planar arm movements with systematically varied joint damping. The model represented a human arm moving in the vertical plane and was controlled by two complementary attractors: one that drove the hand along a circular path and another that stabilized the joints around a nominal configuration [30, 31]. Each attractor included mechanical impedance terms so that the simulated motion reflected realistic interactions between stiffness, damping, and limb dynamics [16, 32–34].

The arm dynamics followed the standard rigid-body formulation,

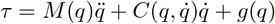

where *τ* ∈ ℝ^2×1^ are the joint torques, 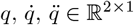 are the joint angles, velocities, and accelerations, *M* (*q*) ∈ ℝ^2×2^is the inertia matrix, 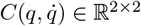 contains Coriolis and centrifugal terms, and *g* (*q*) ∈ ℝ^2×1^ is the gravitational torque vector. Segment lengths, masses, and inertias were chosen to match an average adult male upper and lower arm [35].

Joint torques were generated by combining cartesian hand- and elbow joint-space impedance control:

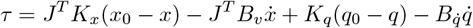

where 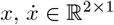 are the hand position and velocity, *J* ∈ ℝ^2×2^ is the Jacobian matrix, *x*_0_ is the circular hand reference trajectory, and *q*_0_ is a fixed postural reference. The impedance matrices were defined as

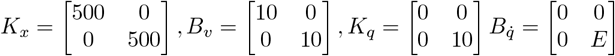

where *K*_*x*_ and *B*_*v*_ are the cartesian hand stiffness and damping matrices, *K*_*q*_ is the joint stiffness matrix, and 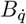 is the joint damping matrix. The scalar parameter *E* ∈ ℝ_≥0_ specifies the damping applied to the elbow joint.

The hand reference trajectory was

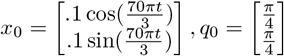

To identify a set of motions that differed primarily in velocity while remaining similar in path shape, we performed a Monte-Carlo search over candidate damping values (0–50 N·m·s/rad), joint stiffness values (0–50 N·m/rad), and trajectory frequencies (0.3–0.7 Hz). The objective was to minimize variance in hand and joint-angle *positions* across conditions while maximizing variance in their *velocities*.

This optimization yielded six trajectories with damping values *E* = 0, 2, 4, 6, 8, 10 N·m·s/rad, joint stiffness fixed at *k* = 10 N·m/rad, and a movement frequency of 0.7 Hz. This configuration yielded small differences in hand paths (SD = 0.0145 m) and elbow angle (SD = 5.52^°^) across the six damping levels, while producing much larger differences in hand tangential velocity (SD = 0.0671 m/s) and elbow angular velocity (SD = 24.8^*°*^/s). These trajectories are shown in Fig. 3. Note that participants viewed only the moving limb during the experiment and did not see the overlaid path traces.

**Fig 3.**
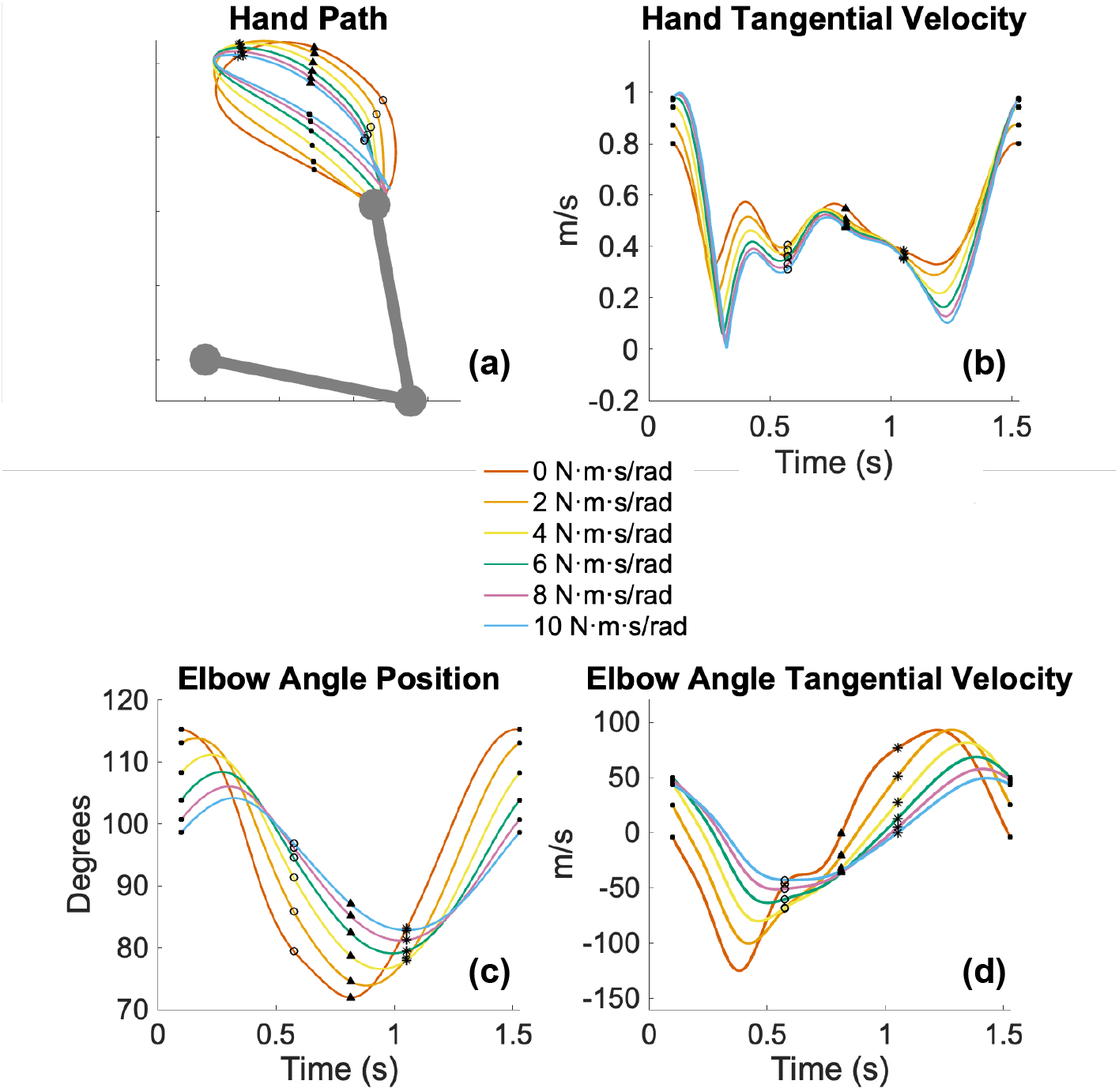
Simulated Hand and Elbow Angle Kinematics. The (a) hand trajectories, (b) hand tangential velocities, (c) elbow angle position, and (d) elbow angle tangential velocities as the damping was varied from 0 to 10 Nms/rad. The markers match the timing between the positions and velocities.

### Task Instruction

All participants were given the following definition of damping: “We define damping as the extent to which an object resists motion in proportion to its velocity. Imagine a door that is hard to pull open quickly–that is because its hinges have high damping. In other words, the door’s highly damped hinges make it harder to move the door at high speeds, causing it to open more slowly.” Prior to the start of the experiment, participants were not presented with examples of “more” and “less” damped arm motions. They also did not receive feedback regarding the accuracy of their estimates at any point during the experiment. Rather, we aimed to assess participants’ innate ability to do this task. Importantly, we intentionally did not provide participants with any details regarding the underlying motion controller.

### Statistical Analyses

To assess whether people can innately perceive different levels of damping (RQ1), we conducted a two-way repeated-measures ANOVA on all participants’ Session 1 ratings with joint damping (6 levels) and block (5 levels) as within-subject factors. Recall a session consisted of 30 trials containing 5 blocks of 6 randomized damping levels.

Additionally, we conducted a three-way repeated-measures ANOVA on Session 2 ratings with coaching group (3 levels) as a between-subjects factor and joint damping (6 levels) and block (5 levels) as within-subject factors.

To determine whether coaching was effective in changing participants’ strategies for visually estimating damping (RQ2), we measured changes from Session 1 to Session 2 in both self-reported and eye-tracking characteristics. Self-reported measures included location (Hand, Elbow, Angle, Other) and motion (Displacement, Velocity, Acceleration, Jerk, Other), while eye-tracking provided additional location measures. These changes were analyzed using a one-way ANOVA with coaching group (3 levels) as a between-subjects factor. We expected Hand Coaching to lead participants to focus on the hand more than the other coaching groups, whereas Angle Coaching was expected to increase attention to the elbow angle compared to the other coaching groups. For both the Hand and Angle Coaching groups, we expected coaching to increase attention to velocity relative to the No Coaching group.

To evaluate whether coaching improved damping perception (RQ3), we fit a linear model relating participants’ damping estimates to the simulated joint damping and extracted the coefficient of determination (*R*^2^) as a performance metric. *R*^*2*^ reflects the proportion of variance in estimates explained by the simulation, with values closer to 1 indicating better performance. We also computed and reported the fitted slope. To test for group differences, we conducted a one-way ANOVA on the change in *R*^*2*^ from Session 1 to Session 2 (and likewise on the change in slope), with coaching group (3 levels) as a between-subjects factor. Post-hoc comparisons were used to test whether the hand and Angle Coaching groups showed greater improvement than the No Coaching group, indicating that coaching significantly enhanced participants’ ability to perceive differences in damping.

Moreover, we tested whether participants’ self-reported use of each motion and location feature predicted performance–quantified as the *R*^*2*^ of their ratings versus simulated damping fits–using a linear mixed-effects model. The location model included reliance on Hand, Elbow, and Angle (percentages) as fixed effects, with a random intercept for subject, while the motion model included reliance on Displacement, Velocity, Acceleration, and Jerk (percentages) as fixed effects, with a random intercept for subject. This allowed us to assess, within a single framework, whether greater reliance on a location or motion characteristic was associated with improved performance.

Finally, we report the NASA TLX survey ratings as changes from Session 1 to Session 2. Each individual loading was submitted to a one-way ANOVA with coaching group (3 levels) as a between-subjects factor.

All analyses were conducted in MATLAB R2022b. Prior to running one-way ANOVAs, data were checked for normality and homogeneity of variance (homoscedasticity) using the Brown–Forsythe test. Unless otherwise noted, all data met these assumptions. However, when the assumption of normality was violated, a Kruskal–Wallis test was used; when the assumption of homoscedasticity was violated, Welch’s ANOVA was used. If both assumptions were violated, Welch’s ANOVA was also used. For all statistical tests, the significance level was set at *α* = 0.05. Where applicable, post-hoc comparisons were adjusted using the Bonferroni correction to control for multiple comparisons.

## Results

Fig. 4 shows all 30 individual participants’ damping estimates plotted against the simulated joint damping values, along with the linear model fit and corresponding *R*^2^. The plots are organized by coaching group, with data from Sessions 1 and 2 overlaid. The greater the *R*^2^, the better the subject’s ability to estimate changes in damping.

**Fig 4.**
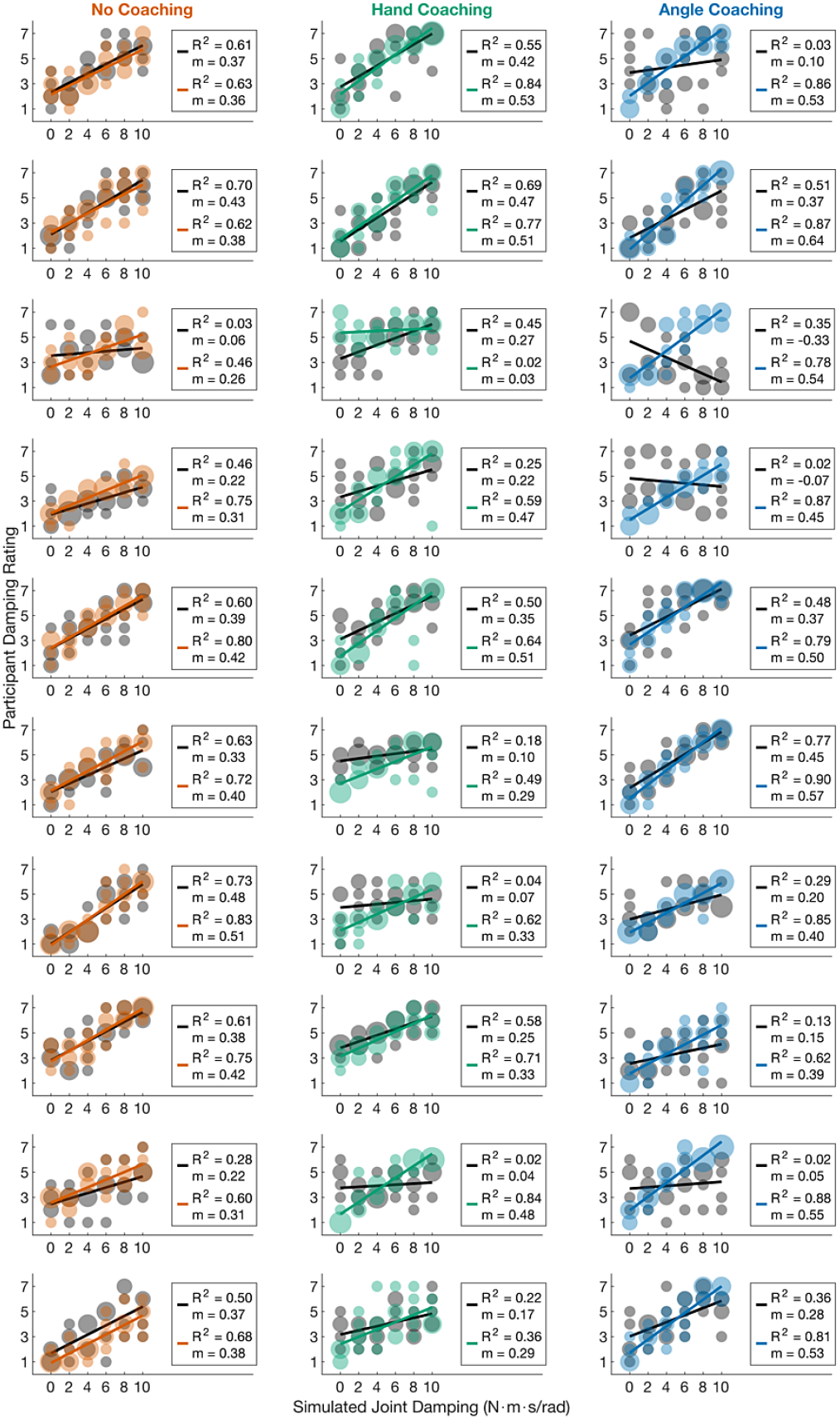
All participants’ damping ratings. Individual participants’ damping rating as a function of simulated damping. The first session ratings are black, while the second session is orange (No Coaching), green (Hand Coaching), and light blue (Angle Coaching). Larger circles represent repeated ratings. *R*^*2*^ and *m* denote the coefficient of determination of the linear fit and the slope, respectively.

Participants across all sessions and coaching groups varied in their ability to estimate damping from motion, as indicated by the overall variation of *R*^*2*^ values (M = 0.54, SD = 0.27) and slopes (M = 0.33, SD = 0.18). Specifically, in Session 1, we found the average *R*^2^ (M = 0.39, SD = 0.24) and slope (M = 0.24, SD = 0.18). Note that for six participants, the linear model fit was not statistically significant (*p >* 0.05; *R*^*2*^ *<* 0.13). Additionally, two participants exhibited negative slopes, although one of these corresponded to a non-significant model fit. In Session 2, Angle Coaching produced the highest average *R*^2^ (M = 0.82, SD = 0.08), followed by No Coaching (M = 0.68, SD = 0.11), and Hand Coaching (M = 0.59, SD = 0.25). For slopes, Angle Coaching again yielded the highest average slope (M = 0.51, SD = 0.08), followed by Hand Coaching (M = 0.38, SD = 0.15), and No Coaching (M = 0.37, SD = 0.07). Only one participant in the Hand Coaching group had an non-significant model fit in Session 2, and no negative slopes were observed.

Although some linear models were non-significant, we included those data in subsequent analyses to avoid biasing our results against participants with inaccurate estimates.

### RQ1. Can humans innately estimate differences in limb damping from visual observation alone when observing movements generated by a simulated two-link arm?

We first asked whether participants could visually discriminate the damping levels of the simulated arm motion without prior coaching. Thus, we performed a two-way repeated-measures ANOVA on ratings with joint damping (6 levels) and block (5 levels) as within-subject factors for all participants’ Session 1 estimates. The analysis revealed a significant main effect of Joint Damping, F(5, 870) = 7.12, p *<*.001, indicating that participants’ ratings varied systematically with the level of damping. In contrast, there was no significant main effect of block, F(4, 870) = 1.83, p =.121, nor was there a significant interaction between Joint Damping and Block F(20, 870) = 1.08, p =.364 (Fig. 5, Black Line).

**Fig 5.**
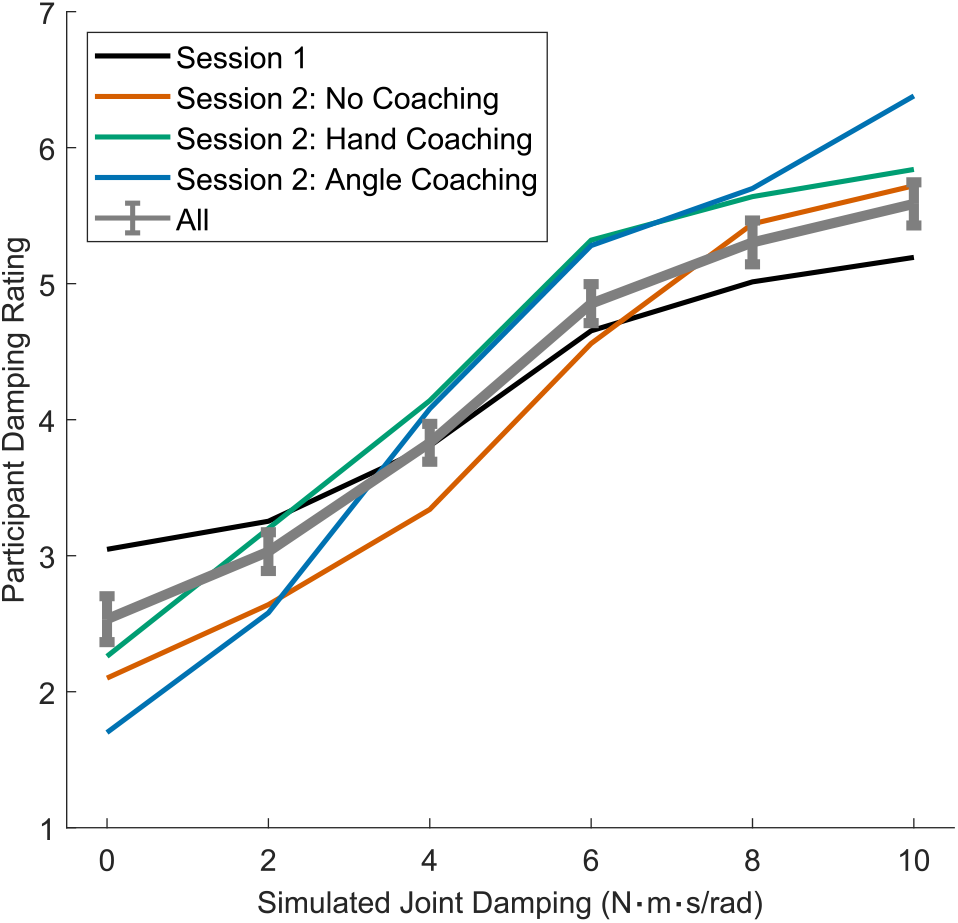
Average damping-estimate results across session and coaching group. Mean perceived damping levels as a function of simulated joint damping are shown for the first session (black) and the second session, separated by coaching group, with No Coaching (orange), Hand Coaching (green), and Angle Coaching (light blue). Increases in accuracy are reflected in a steeper relationship between true and estimated damping. The thick grey line denotes the mean across all sessions; error bars represent 2 SEM.

To assess whether perceptual performance in Session 2 was influenced by coaching, we conducted a three-way repeated-measures ANOVA with coaching group (3 levels) as a between-subjects factor and joint damping (6 levels) and block (5 levels) as within-subject factors. The analysis revealed a significant main effect of Joint Damping, F(5, 870) = 52.50, p *<*.001, indicating that participants’ ratings varied reliably with damping level (Fig. 5). There was also a significant interaction between coaching group and joint damping, F(10, 870) = 3.43, p *<*.001, indicating that the relationship between ratings and damping differed across groups.

No other effects were significant: coaching group, F(2, 397.92) = 1.54, p =.217; block, F(4, 870) = 0.24, p =.914; coaching group × block, F(8, 870) = 0.64, p =.746; joint damping × block, F(20, 870) = 0.51, p =.963; and coaching group × joint damping × block, F(40, 870) = 0.72, p =.906.

Given the significant interaction between Coaching Group and Joint Damping, we next evaluated whether coaching changed the strategies participants used when making these judgments.

### RQ2: Can coaching change the visual attention strategies observers use when estimating damping?

If coaching is effective, it should alter how participants visually sample the motion–either by shifting *where* they look or *which* motion features they rely on. To test this, we compared changes in self-reported focus between Sessions 1 and 2 across the three coaching groups (Figs. 6 and 7). For changes in attention to the hand, a one-way ANOVA revealed a significant main effect of group, F(2, 27) = 5.70, p =.009. Participants in the Hand Coaching group reported the largest increase in focusing on the hand (M = 32.00, SD = 30.8), compared with smaller and negative changes in the No Coaching (M = 4.00, SD = 30.84) and Angle Coaching groups (M = -9.50, SD = 21.40). Post-hoc comparisons showed that Hand Coaching led to a significantly greater increase than Angle Coaching (p =.007) and a marginal increase relative to No Coaching (p =.084), whereas Angle and No Coaching groups did not differ (p =.536).

**Fig 6.**
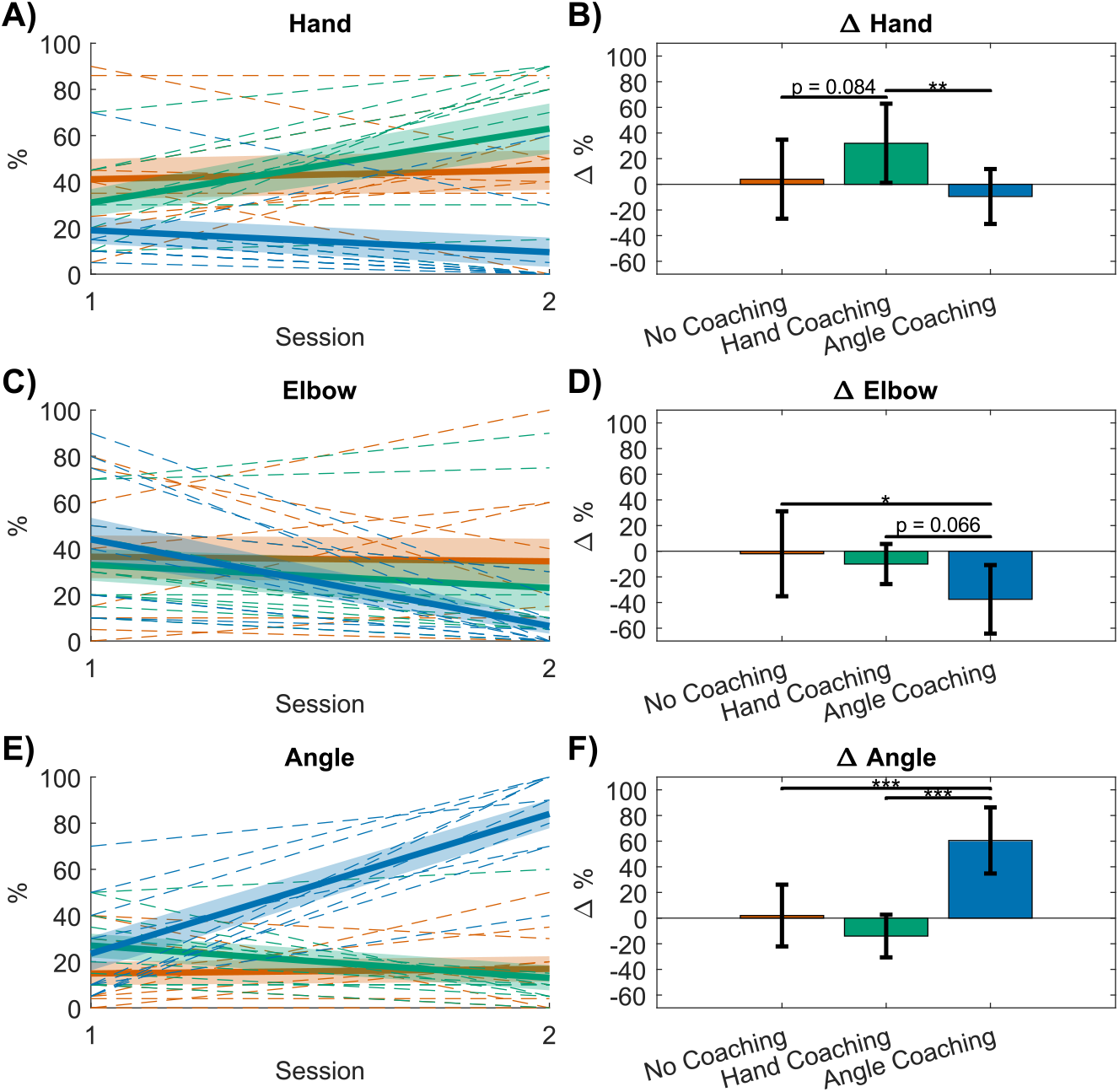
Participants’ self-reported location attention percentages. (A, C, E) The percentage of time participants self-reported looking at the (A) hand, (C) elbow, and (E) angle in the post-session surveys. Thin dashed lines denote individual participants, while the thick solid line and shaded region denote the group average ±1 SEM. Colors indicate coaching groups: No Coaching (orange), Hand Coaching (green), and Angle Coaching (light blue). (B, D, F) The change between Sessions 1 and 2 in the percentage of time participants reported looking at the (B) hand, (D) elbow, and (F) angle, averaged across participants within each coaching group. Error bars represent ± 1 SD. * denotes *P <* 0.05,** denotes *P <* 0.01, and *** denotes *P <* 0.001.

**Fig 7.**
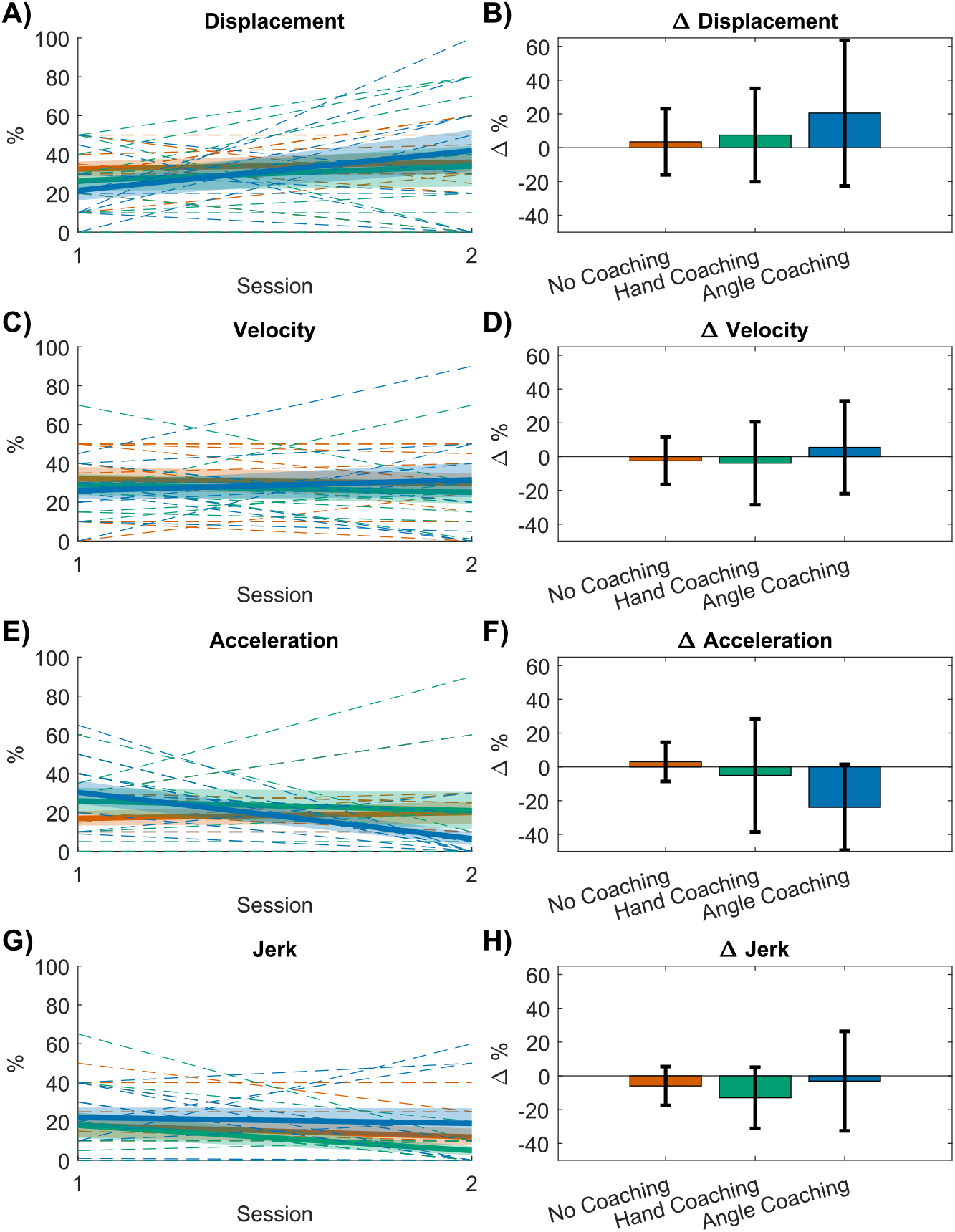
Participants’ self-reported motion attention percentages. (A, C, E, G) The percentage of time participants self-reported looking at the (A) displacement, (C) velocity, (E) acceleration, and (G) jerk in the post-session surveys. Thin dashed lines denote individual participants, while the thick solid line and shaded region denote the group average ± 1 SEM. Colors indicate coaching groups: No Coaching (orange), Hand Coaching (green), and Angle Coaching (light blue). (B, D, F) The change between Sessions 1 and 2 in the percentage of time participants reported looking at the (B) displacement, (D) velocity, (F) acceleration, and (H) jerk, averaged across participants within each coaching group. Error bars represent ± 1 SD.

For changes in attention to the elbow angle, there was a significant main effect of group, F(2, 27) = 16.53, p *<*.001. Participants in the Angle Coaching group reported the greatest increase in focusing on the elbow angle (M = 54.50, SD = 34.11), compared with minimal and negative changes in the No Coaching (M = 8.00, SD = 27.91) and Hand Coaching groups (M = -14.00, SD = 16.63). Post-hoc analyses confirmed that Angle Coaching produced significantly larger increases than both the No Coaching (p =.002) and Hand Coaching groups (p *<*.001), whereas the difference between hand and No Coaching was not significant (p =.186).

For changes in attention to the elbow joint in Cartesian space, the main effect of group was not significant, F(2, 27) = 2.08, p =.144, indicating that participants’ reported focus on the elbow did not differ across coaching conditions.

Together, these results indicate that participants effectively adopted the location-specific visual strategies emphasized in their respective coaching videos–Hand Coaching increased focus on the hand, while Angle Coaching enhanced attention to the elbow angle (Fig. 6).

To validate whether participants accurately self-reported their location-specific viewership, we analyzed the eye-tracking data (See Fig. S5) to measure the average change from Session 1 to Session 2 in location viewing for the three segmentations (Hand, Elbow, Neither). After processing the eye-tracking data for all participants using the method described in the Supporting Information, eight participants had data that passed: No Coaching (2), Hand Coaching (3), Angle Coaching (3). With the limited data, we observed a concurrence in shifts in attention to the hand for Hand Coaching participants, but contrary to survey results, there was no shift in attention to the elbow for Angle Coaching participants. However, given the small amount of reliable data, we refrain from drawing concrete conclusions. Full details are explained in Supporting Information.

For changes in self-reported motion characteristics, no significant differences were found between coaching groups across any of the four categories. Specifically, one-way ANOVAs revealed no main effects of group for displacement, F(2, 27) = 0.22, p =.805; velocity, F(2, 27) = 1.04, p =.366; acceleration, F(2, 27) = 1.14, p =.336; or jerk, F(2, 27) = 0.59, p =.560. These findings suggest that while coaching successfully directed participants’ visual attention to different locations, it did not substantially alter the specific motion features they reported using to estimate damping; participants followed *where* to look (Fig. 6), but not *what* to look for (Fig. 7). These findings raise the question of whether the strategy changes that did occur were responsible for improvements in damping-estimation performance.

### RQ3. Does targeted coaching improve damping-perception accuracy by guiding observers toward informative motion features?

To determine whether coaching improved perceptual performance and identify which strategies drove these improvements, we analyzed changes in estimation accuracy across sessions and examined how strategy use predicted accuracy. We first assessed overall performance changes by evaluating differences in slope and *R*^*2*^ between sessions. We specifically tested whether coaching improved participants’ ability to perceive differences in damping by analyzing the change in model fit (Δ*R*^*2*^) and slope across coaching groups. A one-way ANOVA on Δ*R*^*2*^ revealed a significant effect of coaching group, F(2, 27) = 5.55, p =.010. Participants in the Angle Coaching group showed the largest increase in *R*^*2*^ (M = 0.53, SD = 0.25), followed by the Hand Coaching (M = 0.24, SD = 0.33) and No Coaching (M = 0.17, SD = 0.15) groups. Post-hoc comparisons confirmed that the Angle Coaching group improved significantly more than the No Coaching (p =.011) and Hand Coaching (p =.044) groups, while the hand and No Coaching groups did not differ (p =.822). See Fig. 8 for full results.

**Fig 8.**
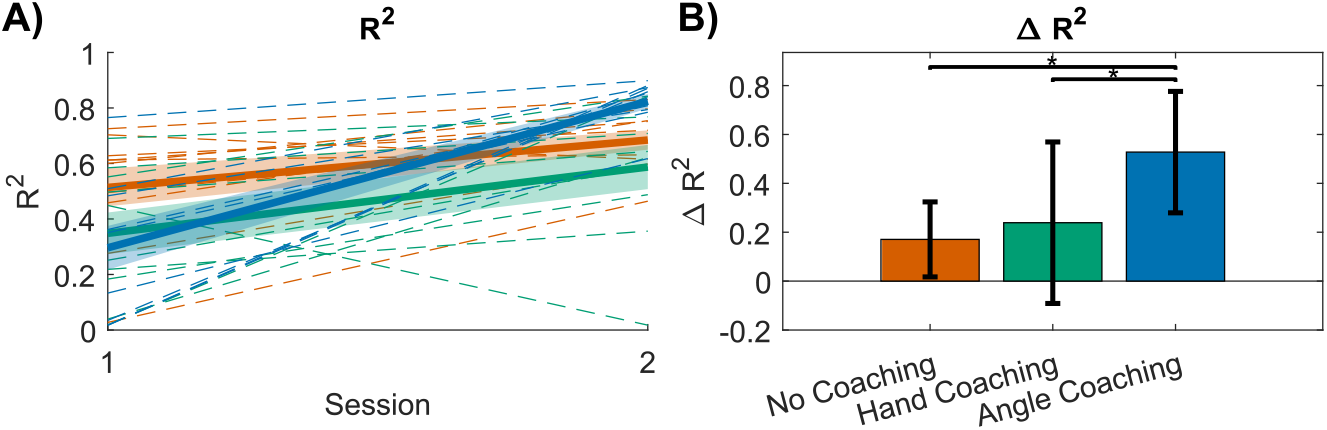
Coefficient of determination (*R*^2^) comparisons across sessions and coaching groups. (A) Individual participants’ *R*^2^ values, obtained from linear fits between the simulated joint damping values and each subject’s damping ratings, are shown for each coaching group in the first and second sessions. Thin dashed lines denote individual participants, while the thick solid line and shaded region denote the group average ±1 SEM. Colors indicate coaching groups: No Coaching (orange), Hand Coaching (green), and Angle Coaching (light blue). (B) The change in *R*^2^ between Session 1 and Session 2, averaged across participants, is shown for each coaching group. Error bars represent ± 1 SD. * denotes *P <* 0.05 For slope, there was a violation of normality (p =.037), and the Kruskal-Wallis test revealed a significant main effect of group, *χ*^2^(2) = 15.11, p *<*.001. The Angle Coaching group again showed the greatest increase in slope (M = 0.35, SD = 0.23), compared to the Hand Coaching (M = 0.14, SD =.18) and No Coaching groups (M = 0.05, SD = 0.07). Post-hoc analyses revealed that the Angle Coaching group exhibited significantly greater improvement than the No Coaching group (p =.003), whereas there were no differences between the hand and Angle Coaching (p =.117) and hand and No Coaching groups (p =.137). See Fig. 9 for full results.

To test whether specific strategy changes accounted for these improvements, we next used a linear mixed-effects (LME) model relating performance to participants’ self-reported use of each motion feature. Specifically, we fit separate LME models for location and motion features, with *R*^*2*^ from participants’ damping-estimation fits as the dependent variable and a random intercept for subject.

For the location model (Fig. 10), greater reported focus on the hand (*β* = 0.0046 ± 0.0019, t(55) = 2.34, p =.023) and elbow angle (*β* = 0.0048 ± 0.0019, t(55) = 2.48, p =.016) were both significant positive predictors of performance, whereas focus on the elbow was not (*β* = 0.0017 ± 0.0019, p =.371).

For the motion model (Fig. 11), reliance on acceleration was a significant negative predictor of performance (*β* = -0.0052 ± 0.0020, t(54) = -2.60, p =.012), suggesting that participants who attended more to acceleration performed worse. No significant effects were found for displacement (p =.95), velocity (p =.11), or jerk (p =.90).

### NASA Task Load Index

Finally, we examined whether coaching affected subjective workload. To evaluate differences in perceived workload across coaching groups, we analyzed changes in NASA TLX ratings from Session 1 to Session 2 using one-way ANOVAs for each subscale (Fig. 12).

For mental demand, a one-way ANOVA revealed a significant main effect of group, F(2, 27) = 4.13, p =.027. Participants in the Angle Coaching group reported the greatest reduction in mental demand (M = -0.30, SD = 0.48), followed by the Hand Coaching group (M = 0.10, SD = 0.99) and the No Coaching group (M = 1.00, SD = 1.41). Post-hoc comparisons indicated that perceived mental demand decreased significantly more for the Angle Coaching group than for the No Coaching group (p =.024), while differences between No Coaching and Hand Coaching (p =.147) and between Hand Coaching and Angle Coaching (p =.668) were not significant.

For physical demand, a violation in normality (p =.010) and a violation in homoscedasticity (p =.024) were observed, thus a Welch’s ANOVA was conducted, which found an insignificant main effect of group, F(2, 14.96) = 0.87, p =.440.

For frustration, a violation of normality was observed (p =.013). A Kruskal-Wallis test confirmed no group differences, *χ*^2^(2) = 2.03, p =.363.

No significant main effects of group were observed for Temporal Demand, F(2, 27) = 0.84, p =.445; Effort, F(2, 27) = 1.02, p =.376; and Performance, F(2, 27) = 1.45, p =.253.

## Discussion

In this study, we investigated whether humans can visually perceive differences in limb damping from motion alone and whether targeted coaching can enhance this ability. Using abstract two-link arm simulations with systematically varied joint damping, participants completed two blocks of rating tasks separated by a brief coaching intervention. Across sessions, we combined subjective reports, eye-tracking, and performance metrics (*R*^2^ and slope of rating–damping fits) to assess not only participants’ innate perceptual abilities but also how coaching affected the visual strategies they used.

Humans can innately perceive differences in damping solely from their visual observation of motion. Even in Session 1, before any instructional feedback was given, participants’ estimates scaled monotonically with the true damping level of the simulated arm (Fig. 5) (Black Line). This is remarkable because damping is defined as a ratio of two variables, velocity *and* force in the formula, *F* = *bv* [36]. Despite having no prior training in how to visually estimate damping, participants produced systematic and reliable judgments, suggesting that they may be drawing on internal predictive models shaped by their own motor experience and the shared neural resources underlying action and perception [9–11]. Taken together, these results demonstrate that humans possess an innate ability to extract a hidden component of mechanical impedance from vision alone, extending previous findings on visual stiffness perception [5–7].

Coaching effectively shaped participants’ strategies. However, its influence was specific: coaching reliably changed *where* participants looked, but not *what* motion features they reported using. We observed a significant increase in self-reported attention to the hand for participants in the Hand Coaching group, and a significant increase in attention to the elbow angle for participants in the Angle Coaching group. These shifts confirm that participants adopted the coached visual locations.

While the coaching video shown to participants in the Hand Coaching group specifically recommended that they attend to the *velocity* of the hand’s motion (See S1) and the coaching video shown to participants in the Angle Coaching group recommended attending to the *velocity* of the opening and closing of the elbow joint (See S2), participants in all groups had statistically-insignificant changes in self-reported motion strategy (Fig. 7). Moreover, self-reported use of displacement, velocity, or jerk did not significantly predict rating performance, whereas reduced attention to acceleration was associated with significantly better accuracy (Fig. 11). Together, these findings indicate that participants did not adopt the motion-feature instructions emphasized in the coaching videos. Instead, participants may have relied on more informative cues than those highlighted in the coaching materials. Overall, these results help identify which forms of coaching are most useful for perceptual learning and emphasize the importance of training paradigms that explicitly teach observers to detect the motion features that truly reveal latent dynamic properties.

Although these results contradict our original hypothesis, they are consistent with prior work. Previous research has shown that motion features containing path information are more intuitive for estimating joint stiffness compared to features based on temporal information [5]. Studies have also found that humans are better at perceiving motion differences based on the spatial path of objects, independent of the temporal frequency that determines velocity [37]. Thus, participants’ tendency to rely on spatial rather than temporal motion cues aligns with established perceptual biases and may explain why the velocity-focused instructions were not adopted.

Moreover, as seen in Fig. 3, the hand path distinctly flattens as damping increases, whereas the corresponding variation in hand velocity is far less pronounced. Because the Hand Coaching video presented a side-by-side comparison of hand paths in low- and high-damping simulations (See S1), participants in this group likely drew on these displacement differences when forming their strategies in Session 2. This is supported by several remarks from Hand Coaching participants in their Session 2 self-reported survey, including, “focus was on parabolic trajectory of end point as I was tracing the point path … when the parabola was very small it was high damping but when it was making a larger round it was low damping”, “size of the loop”, “areas drawn by the hand and compare their sizes with more circular shapes indicating higher velocities and more elongated ones slower”, “path traced by the Hand.” This reliance on spatial path information is consistent with participants’ general tendency to favor displacement cues over velocity-based features.

Coaching improved participants’ performance in estimating damping. Across all three coaching groups, participants improved their *R*^2^ and slope from Session 1 to Session 2. (Fig. 8 and 9). Participants in the No Coaching group may have improved simply due to increased familiarity with the range of possible simulations after completing Session 1. Additionally, one participant in the No Coaching group pointed out how seeing the diagram with labeled sections for “Hand”, “Elbow”, and “Angle” in the survey following Session 1 helped them recognize the value of directing attention to one of those specific regions of the arm when making their judgments in Session 2 (Fig. 2). These incidental sources of learning may partially explain the modest improvements observed in the No Coaching group.

**Fig 9.**
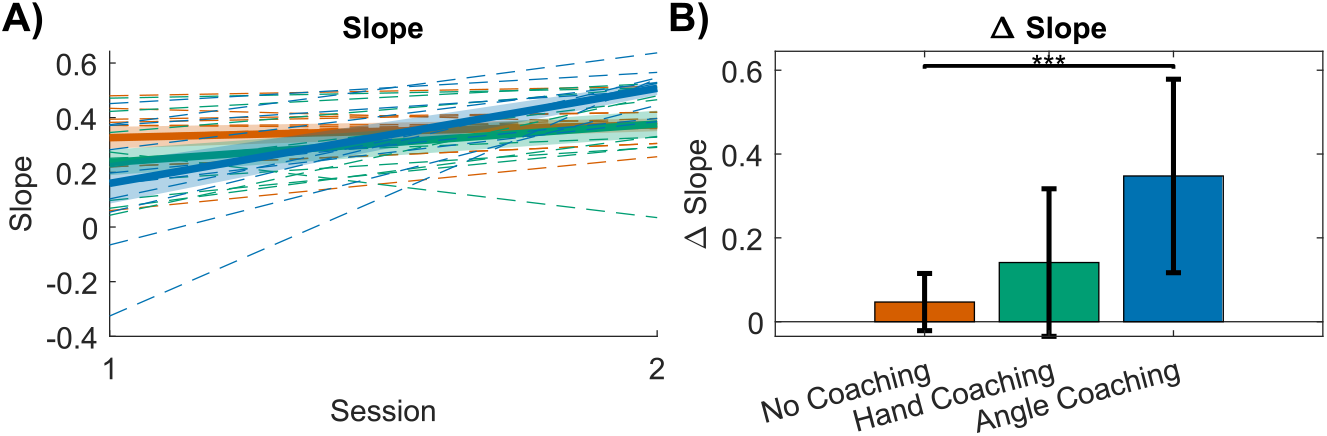
Slope comparisons across sessions and coaching groups. (A) Individual participants’ slope values, obtained from linear fits between the simulated joint damping values and each subject’s damping ratings, are shown for each coaching group in the first and second sessions. Thin dashed lines denote individual participants, while the thick solid line and shaded region denote the group average ±1 SEM. Colors indicate coaching groups: No Coaching (orange), Hand Coaching (green), and Angle Coaching (light blue). (B) The change in slope between Session 1 and Session 2, averaged across participants, is shown for each coaching group. Error bars represent ± 1 SD. *** denotes *P <* 0.001

**Fig 10.**
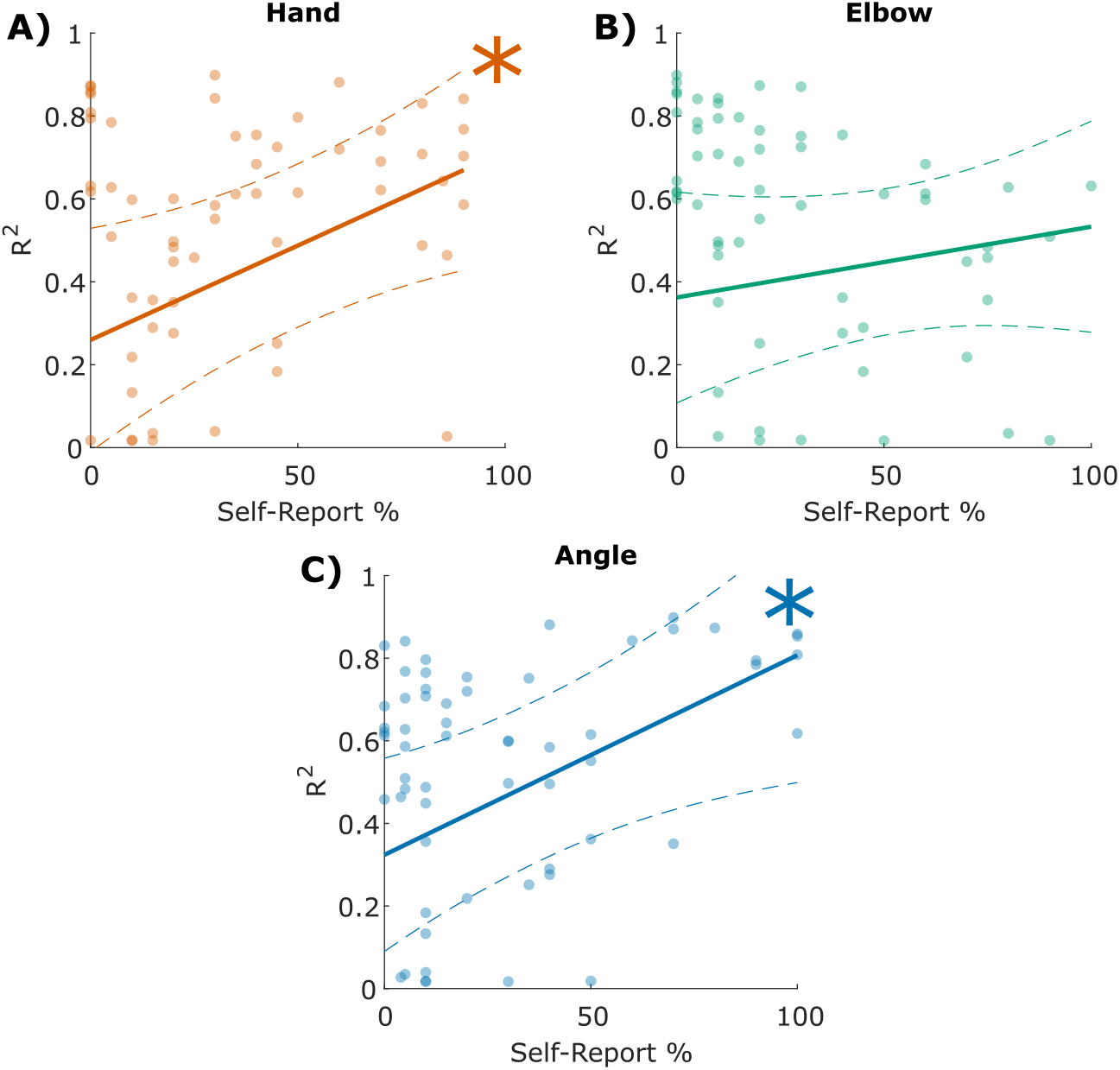
Relationship between self-reported location feature use and damping-estimation accuracy. Linear mixed-effects model fits are shown for the association between participants’ *R*^*2*^ values and the percent of time they reported using the (A) hand, (B) elbow, and (C) angle when making their judgments. Points denote individual participants, solid lines indicate the fitted fixed-effect regression for each predictor (with the other predictors held at their median values), and dashed lines show the 95% confidence band. Asterisks denote significant fixed-effect coefficients (*p <* 0.05).

**Fig 11.**
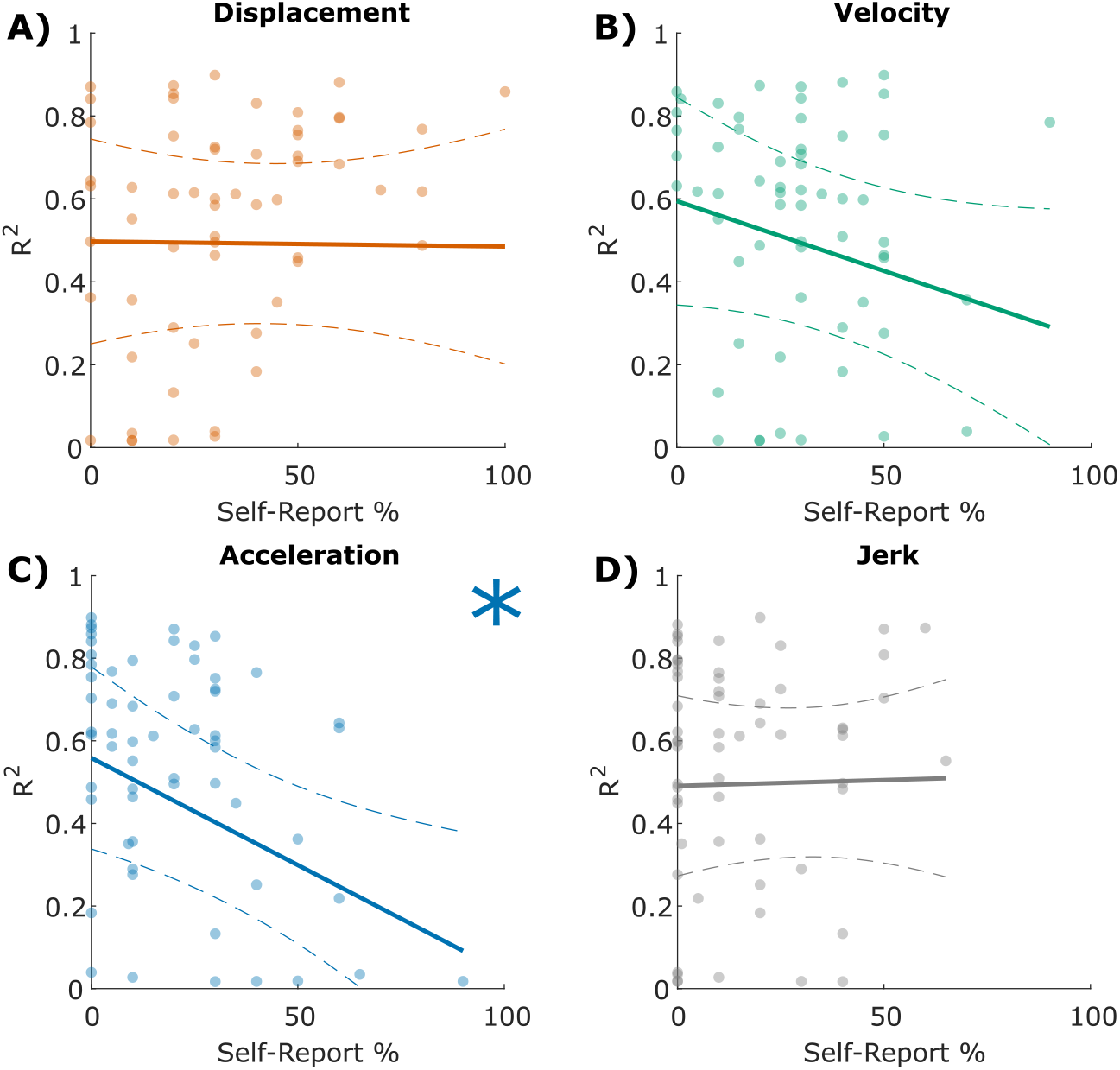
Relationship between self-reported motion feature use and damping-estimation accuracy. Linear mixed-effects model fits are shown for the association between participants’ *R*^2^ values and the percent of time they reported using the (A) displacement, (B) velocity, (C) acceleration, and (D) jerk when making their judgments. Points denote individual participants, solid lines indicate the fitted fixed-effect regression for each predictor (with the other predictors held at their median values), and dashed lines show the 95% confidence band. Asterisks denote significant fixed-effect coefficients (*p <* 0.05).

**Fig 12.**
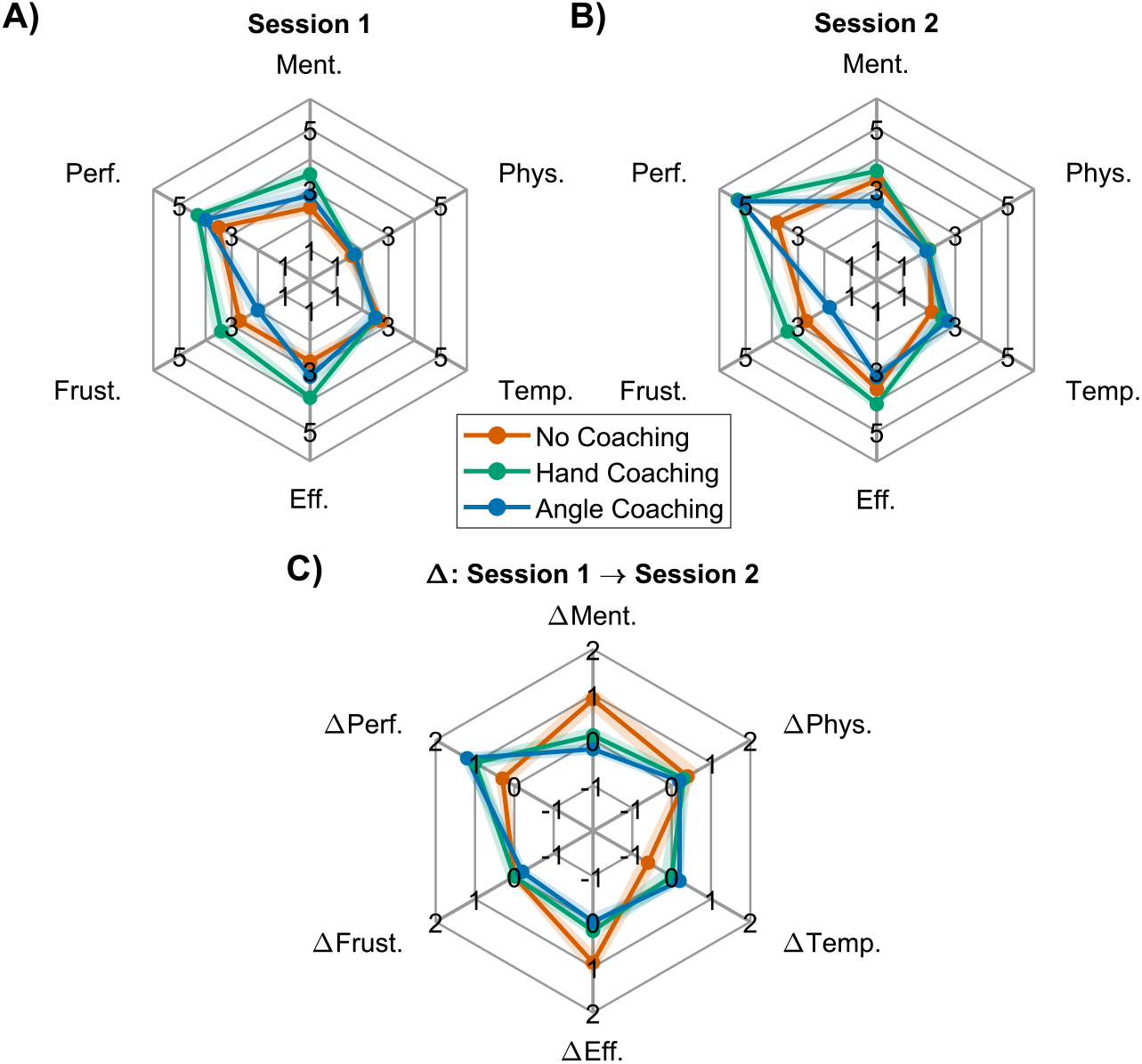
NASA-TLX workload ratings. NASA-TLX ratings for (A) Session 1, (B) Session 2, and (C) the change in each workload dimension from Session 1 to Session 2 are shown for each coaching group: No Coaching (orange), Hand Coaching (green), and Angle Coaching (light blue). Thick lines denote group averages and shaded regions represent ± 1 SEM. Ment., Phys., Temp., Eff., Frust., and Perf. correspond to mental demand, physical demand, temporal demand, effort, frustration, and performance, respectively.

The LME location effects model found that increased attention to the hand or angle resulted in significantly better performance (Fig. 10). This aligns with the coaching strategies we provided that suggested where participants should focus their attention to more accurately distinguish damping levels.

Angle Coaching was most effective. Participants in the Angle Coaching group experienced the highest and most statistically-significant increase in *R*^2^ and slope from Session 1 to Session 2 compared to the other coaching groups (Fig. 8 and 9). This makes sense because the manipulated parameter in the simulations was the damping applied at the elbow joint, making elbow-angle motion the most informative cue for distinguishing damping levels. Moreover, while units were not necessarily comparable, the elbow trajectory exhibited greater variance than the hand trajectory, providing a richer signal for perceptual discrimination. This finding aligns with results from a previous study on visual stiffness perception, in which participants tended to rely on joint-space information when estimating stiffness [5].

There was only one participant out of the 30 who did not improve from Session 1 to Session 2. This participant was in the Hand Coaching group (Fig. 4: column 2, row 3); according to their self-reported strategy for Session 2, they did not follow the instructed strategy of attending to hand velocity. Instead, they reported that they “focus[ed] on the acceleration of the elbow joint - mainly on the bottom right turn.” This is notable because increased attention to acceleration was associated with poorer performance in the group-level LME analysis (Fig. 11), suggesting that this participant adopted a particularly ineffective strategy.

Finally, the NASA TLX results suggest that coaching primarily reduced participants’ perceived mental demand, particularly for those who received Angle Coaching, while other aspects of perceived workload did not differ significantly across coaching groups (Fig. 12). This reduction in mental demand for the Angle Coaching group is consistent with the fact that angle information was the most diagnostic cue for distinguishing damping levels. By directing participants to the most informative feature, coaching may have reduced the cognitive effort required to search the motion for relevant cues, allowing participants to adopt a more efficient and less demanding perceptual strategy. More broadly, these results suggest that effective coaching not only improves perceptual accuracy but also lowers the cognitive burden of the task, an important consideration for designing training paradigms in domains where sustained perceptual judgment is required.

### Limitations

Due to the nature of the actual simulations being spread across 6 damping levels while participants provided ratings on a 1–7 Likert scale, we used the *R*^2^ linear fit statistic rather than the root mean squared error (RMSE). We choose a scale of 1-7 to maintain methodological consistency and comparability with the prior stiffness studies [5–7]. Moreover, we used the slope of the linear fit as an additional performance statistic that we calculated for each participant, where values closer to 1 indicate stronger sensitivity to changes in damping. Future work could explore study designs that use RMSE as an accuracy metric or incorporate alternative modeling approaches, such as nonlinear fits, to better characterize perceptual performance across the discrete damping levels.

A limitation of using eye-tracking to validate the accuracy of participants’ self-reported location strategies is that although the survey asked participants to estimate the time they were focused on the elbow point versus the elbow angle, the low spatial accuracy of the eye-tracking system made it difficult to reliably distinguish these regions. We intended to use eye-tracking to validate participants’ reported strategies, but device limitations prevented meaningful differentiation. One likely reason is that participants moved their heads during the session, causing the glasses to shift position and degrade the calibration over time. Future work could mitigate this issue by using more stable or higher-precision eye-tracking systems, incorporating head-position compensation, or designing tasks that constrain head movement or allow for frequent recalibration.

### Impact

Humans possess a remarkable capacity to learn, adapt, and generalize motor behaviors across contexts [38–44]. This learning is not restricted to one’s own actions; observations of others also shape internal models of motor behavior [45], reflecting the shared neural resources of perception and action. Neuroimaging evidence shows that a mirror-neuron system engages during both movement execution and action observation [9, 10], supporting the idea that humans interpret observed motion by mapping it onto their own motor circuitry [11]. Yet despite this powerful perceptual–motor coupling, the mechanistic variables that generate movement (e.g., muscle activation, neuromotor commands, and limb impedance) are normally hidden from view [46]. Our findings contribute to a growing body of evidence that humans can nevertheless infer latent dynamic properties from visual motion. Prior work demonstrates that people can estimate stiffness from visual observation alone [5], and the present study extends this capacity to damping, a velocity-dependent component of impedance. This perceptual sensitivity may arise because the simulated controller resembles biologically inspired control strategies humans use when generating their own movements, enabling observers to draw upon internal predictive models when interpreting visual motion [9, 10]. The insight that humans can visually access both stiffness and damping aligns with longstanding theories that impedance regulation is central to human motor control and learning [14–18, 47, 48], suggesting that impedance may be a perceptually privileged variable embedded within observers’ internal models.

Beyond demonstrating that damping is visually accessible, the present study shows that the strategies used to perceive impedance are malleable: participants improved their damping-estimation accuracy after targeted coaching. This finding resonates with behavioral and neurological evidence showing that humans, including infants, adults, and non-human primates, leverage observation to imitate motor behaviors [1, 12, 13, 49–53]. Our results suggest that part of this capacity may stem from the ability to infer latent dynamic properties, not just kinematics, from others’ movements. The result that coaching selectively enhanced these perceptual strategies demonstrates that the underlying perceptual models are malleable. Together, these findings clarify which features of motor behavior can be gleaned through visual observation and strengthen the view that humans rely on internal, impedance-based representations when perceiving and predicting movement. As such, this work advances fundamental understanding of perception–action coupling, and the dynamic variables the nervous system encodes during action interpretation.

This ability to infer and refine visual perception of latent limb dynamics is particularly consequential in clinical settings, where such judgments directly inform diagnosis and treatment. For example, rigidity and spasticity are displacement- and velocity-dependent abnormalities in muscle tone that significantly impair the quality of life of stroke survivors. These impairments are still assessed primarily through hands-on examinations in which physical therapists rely on subjective judgments of resistance during passive movement [28]. This reliance on subjective evaluation leads to substantial variability across clinicians [28]. Moreover, rigidity and spasticity map directly onto the mechanical properties of stiffness and damping, respectively, and both our study and prior work show that these properties can be inferred from the visual observation of motion alone. By identifying the visual cues humans naturally use to estimate stiffness and damping and demonstrating that these cues can be strengthened through targeted coaching, this work moves toward physics-based, quantitative markers of rigidity and spasticity that clinicians can apply at the bedside. Grounding these assessments in mechanical metrics such as stiffness and damping can make clinical practice more consistent, quantitative, and reproducible. In addition, the perceptual principles revealed here can inform the design of digital coaching tools that train clinicians where to focus their attention and which motion features are most diagnostically informative when evaluating limb impedance. By reducing the learning curve and improving the consistency of clinical judgments, such tools have the potential to expand the number of therapists capable of delivering high-quality stroke rehabilitation and promote more reliable and widespread patient care [54].

Beyond its implications for clinician training, this work also points toward the development of automated, video-based tools for remotely monitoring stroke recovery [55]. A computer system that analyzes a video of a patient moving their limbs can provide objective measurements of stiffness (rigidity) and damping (spasticity) without requiring an in-person clinic visit. Modern pose-estimation methods such as OpenPose [56], MediaPipe [57], and DeepLabCut [58] make it possible to extract joint kinematics from everyday video captured on a phone, laptop, or tablet, enabling movement data to be collected outside traditional clinical environments. Such technology is particularly valuable in rural and underserved regions, where the availability of licensed physical therapists is much lower and access to specialized care is limited [59]. Integrating computer vision, physics-based models, and machine learning can therefore support more accessible, affordable, and consistent telehealth rehabilitation. By supplementing in-person care with reliable remote assessments, these tools have the potential to improve continuity of treatment and expand high-quality rehabilitation services to patients who might otherwise receive little or no follow-up care.

Additionally, robot-assisted minimally invasive surgery (RMIS) requires surgeons to infer tissue stiffness and other mechanical properties from vision alone, as most systems provide limited or no haptic feedback [60, 61]. Over time, expert surgeons develop internal perceptual models that allow them to interpret subtle deformation cues and anticipate tissue behavior, but acquiring these skills demands extensive practice [62–64]. Evidence shows that experts significantly outperform novices on tasks that rely on visually inferring tactile properties. For example, accurately judging membrane thickness in da Vinci simulations [65]. Our findings suggest that structured coaching may accelerate the development of these internal models. Just as targeted instruction improved participants’ ability to visually estimate damping in an abstract limb simulation, similar strategies could help surgical trainees focus on the most informative visual cues during tissue manipulation. By explicitly guiding where and how to allocate visual attention, coaching-driven approaches may shorten training times, standardize perceptual skills, and ultimately support safer, more precise surgical decision-making [66].

Taken together, our results show that damping, like stiffness, can be visually accessed and systematically improved through training. Thus, mechanical impedance emerges as a core variable in perception–action coupling with clear implications for motor control, rehabilitation, telehealth, and surgical expertise.

## Conclusion

We showed that humans can correctly infer changes in limb damping from visual perception alone. This result was especially robust for participants who received coaching to focus on the opening and closing of the elbow joint angle. Moreover, these observations reinforce existing evidence that humans rely more on path information than velocity when estimating limb impedances from motion. Our findings also support the idea that humans’ ability to visually recognize changes in stiffness and damping may arise from the congruence between the simulated controller used to generate the movements and the biological control strategies people rely on when interpreting motion. Finally, our coaching insights support the belief that training people to do a task beyond their innate ability will have meaningful implications in clinical settings, for example, for training physical therapists to better diagnose patients with spasticity, as well as for training surgeons to better perform procedures with robots lacking haptic feedback. Together, these results demonstrate that even deeply hidden dynamic properties can be revealed through vision and shaped through coaching, opening new opportunities to design training that strengthens human perceptual models in both science and clinical practice.

## Data Availability

All subject data, produced simulations, and initial analysis are available on GitHub at link: https://github.com/michaelwestjr/Huang_2026_PerceivingLatentDynamics-InnateAndCoachableVisualEstimationOfLimbDamping

