## Supplemental information for "Perceiving latent dynamics: Innate and coachable visual estimation of limb damping"

### Supporting information

**S1. Hand Coaching Video** This two-minute clip was shown to the ten participants in the Hand Coaching group after completing Session 1 and before Session 2. The video highlights changes in the speed of the hand path as an effective strategy for determining damping.

**S2. Angle Coaching Video** This two-minute clip was shown to the ten participants in the Angle Coaching group after completing Session 1 and before Session 2. The video highlights changes in the opening and closing speed of the elbow angle as an effective strategy for determining damping.

**S3. Experiment Demo Video** This is the exact graphical user interface that a participant navigated to begin viewing a damped two-link arm simulation, skip as needed, and rate their perceived damping level.

#### Eye-Tracking

##### Eye-Tracking Methods

In an attempt to provide quantitative validation of the location strategies described by participants in the self-reported survey, participants wore eye-tracking glasses (Pupil Core, Berlin) to monitor their gaze as they watched the simulations. We used a Python and MATLAB pipeline to directly translate participants' gaze position into comprehensive statistics on the percentage of time they looked at specific components of the two-link arm.

The eye-tracking glasses contain three cameras: two that closely record the wearer's eyes to determine gaze through pupil detection, and one that faces outward to capture the computer screen environment from the wearer's point of view ("world view").

Prior to data collection, the eye-tracking glasses were calibrated to account for gaze offset. At the beginning of each session, the MATLAB user interface displayed a small white dot at the center of a black screen for five seconds, followed by the same dot appearing sequentially for five seconds at each of the four corners of the square region approximately where the two-link arm simulation would later be shown. Participants were instructed to fixate on the white dot at each location without moving their head.

MATLAB's DLTdv8a (data video version 8a) app was used to dynamically track the x and y coordinates of the calibration dots from the participant's "world view" video and compared these values to the matching gaze coordinates for each frame. Specifically, the *mean* calibration dot coordinate and the *median* gaze coordinate at each of the four corners was used to compute a bilinear interpolation that mapped offset gaze points to their true screen locations. The "world view" was decomposed into individual frames, and a contrast-based computer vision program implemented with the OpenCV Python library was applied to each frame to detect the hand and elbow positions of the two-link arm simulation displayed on screen. After transforming the hand and elbow coordinates into the same coordinate system as the gaze data and applying the calibration-based interpolation to correct for gaze offset, each frame was classified as showing the participant looking at the hand, the elbow, or neither. A frame was classified as *hand* if the gaze point fell within a radius equal to half the forearm length (from the hand to the elbow joint), and classified as *elbow* if the gaze point fell within a radius equal to half the forearm length from the elbow to the hand (Fig. S4). If the gaze point did not fall within either radius, the frame was classified as *neither* (Fig. S4). This classification was performed frame by frame across the entire session. Additionally, any frame with a gaze-confidence value below 0.85 was discarded. Finally, the percentage of time each participant spent looking at the hand, the elbow, or neither across all 30 trials of the session was aggregated. If a participant's tracking report indicated looking at "Neither" for $\geq 50\%$ of the time for a single session, both sessions for that participant would be discarded.

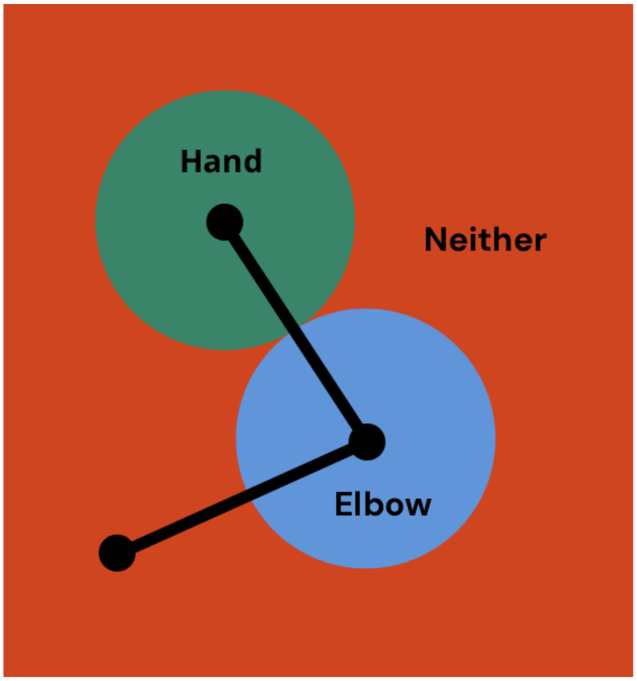


**Fig S4. Gaze classification.** The gaze at a single frame would be classified in the Neither, Hand, or Elbow category based on its location in relation to the arm simulation.

##### Eye-Tracking Results

For shifts in attention to the hand, the results match the expectation that the Hand Coaching group would show the largest increase (M = 49.22), as well as a slight decrease for the Angle Coaching group (M = -5.83). Interestingly, the two No Coaching participants also shifted over to increased attention to the hand in Session 2 compared to Session 1 (M = 38.72). Comparing these eight participants with their self-reported strategies, the Hand Coaching participants had an average increase in focusing on the hand (M = 40), the No Coaching participants also had an average increase in focusing on the hand (M = 35), and the Angle Coaching participants had a slight decrease in focusing on the hand (M = -10). These self-reports generally match the eye-tracking results.

As for the attention to the elbow, there were decreases in viewership for the participants in the No Coaching (M = -40.13) and Hand Coaching (M = -40.83) groups. There was also a slight decrease for the Angle Coaching group (M = -7.51), contrary to our original hypothesis. This is paired with a slight increase in tracked data for "Neither", which suggests that the eye-tracking data for these participants were likely inaccurate, thus producing unexpected results. Comparing these participants with their self-reported strategies, the Angle Coaching participants self-reported a large increase in focusing on the angle (M = 58.33), while No Coaching participants self-reported a slight increase (M = 17.5) and Hand Coaching participants self-reported a decrease (M = -13.33) (See Fig S5.).

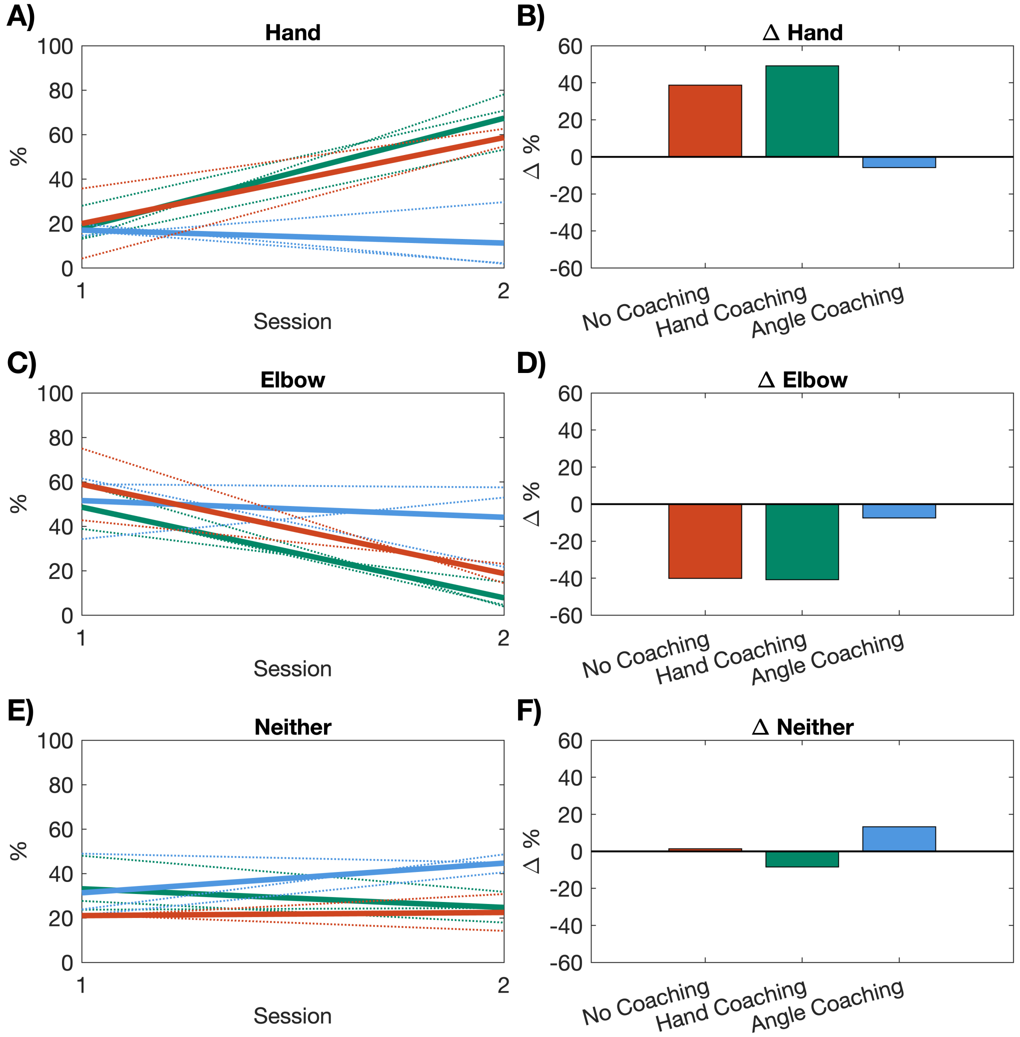


**Fig S5. Participants' eye-tracking location percentages.** This figure was generated with 8 of the 30 participants' data. (A, C, E) The percentage of time participants were tracked looking at the (A) hand, (B) elbow, or (C) neither. Thin dashed lines denote individual participants, while the thick solid line denotes the group average. Colors indicate coaching groups: No Coaching (orange), Hand Coaching (green), and Angle Coaching (light blue). (B, D, F) The change between Sessions 1 and 2 in the percentage of time participants were tracked looking at the (B) hand, (D) elbow, or (F) neither, averaged across participants within each coaching group.
